# PLANT UNCOUPLING MITOCHONDRIAL PROTEIN 2 localizes to the Golgi

**DOI:** 10.1101/2023.05.06.539686

**Authors:** Philippe Fuchs, Elisenda Feixes-Prats, Paulo Arruda, Elias Feitosa-Araújo, Alisdair R. Fernie, Christopher Grefen, Sophie Lichtenauer, Nicole Linka, Ivan de Godoy Maia, Andreas J. Meyer, Sören Schilasky, Lee J. Sweetlove, Stefanie Wege, Andreas P. Weber, A. Harvey Millar, Olivier Keech, Igor Florez Sarasa, Pedro Barreto, Markus Schwarzländer

**Affiliations:** Institute of Plant Biology and Biotechnology (IBBP), Universität Münster, Schlossplatz 8, D-48143 Münster, Germany; Institute of Crop Science and Resource Conservation (INRES), Rheinische Friedrich-Wilhelms-Universität Bonn, Friedrich-Ebert-Allee 144, D-53113 Bonn, Germany; Centre for Research in Agricultural Genomics (CRAG), Campus UAB Bellaterra, 08193 Barcelona, Spain; Genomics for Climate Change Research Center, Universidade Estadual de Campinas, 13083-875 Campinas, Brazil; Department of Molecular Physiology, Max Planck Institute of Molecular Plant Physiology, D-14476 Postdam-Golm, Germany; Institute of Molecular and Cellular Botany, Ruhr-Universität Bochum, Universitätsstrasse 150, D-44780 Bochum, Germany; Institute of Plant Biochemistry, Cluster of Excellence on Plant Science (CEPLAS), Heinrich-Heine University Düsseldorf, D-40225 Düsseldorf, Germany; Institute of Biosciences, São Paulo State University (UNESP), 18618-970 Botucatu, Brazil; Department of Biology, South Parks Road, University of Oxford, OX1 3RB Oxford, United Kingdom; ARC Centre of Excellence in Plant Energy Biology, School of Molecular Sciences, The University of Western Australia, 6009 Perth, Australia; Department of Plant Physiology, Umeå Plant Science Centre, Umeå University, S-90187 Umea, Sweden; Institut de Recerca I Tecnología Agroalimentàries (IRTA), Edifici CRAG, Barcelona, Spain

## Abstract

Mitochondria act as cellular hubs of energy transformation and metabolite conversion in most eukaryotes. Plant mitochondrial electron transport chains are particularly flexible, featuring alternative components, such as ALTERNATIVE NAD(P)H DEHYDROGENASES and ALTERNATIVE OXIDASES (AOXs), that can bypass proton translocation steps. PLANT UNCOUPLING MITOCHONDRIAL PROTEINS (named PUMPs or plant UCPs) have been identified in plants as homologues of mammalian Uncoupling Proteins (UCPs), and their biochemical and physiological roles have been investigated in the context of mitochondrial energy metabolism. To dissect UCP function in Arabidopsis, the two most conserved (UCP1 and UCP2) have been targeted in recent work by combining mutant lines to circumvent potential functional redundancy *in vivo*. Such approaches rely on the assumption that both proteins reside in the inner mitochondrial membrane as a prerequisite for functional redundancy. Yet, contradicting results have been reported on UCP2 localization in plants. Here we provide evidence that, conversely to UCP1, which is an abundant inner mitochondrial membrane protein, UCP2 localizes to the Golgi rather than to mitochondria. Based on multiple lines of new and prior evidence, we summarize the consensus view that we have reached and provide an example of how open, critical exchange within the research community is able to constructively address ambiguities. Our observations and considerations provide direction to the ongoing discussion about the functions of UCP proteins. They further offer new perspectives for the study of Golgi membrane transport and subcellular targeting principles of membrane proteins. Since 20 to 30 % of genes in plant genomes are predicted to encode transmembrane proteins and the function of most of those proteins has not been experimentally investigated, we highlight the importance of using independent evidence for localization as a prerequisite for understanding physiological function of membrane proteins.

---

Uncoupling Proteins (UCPs) are integral components of mitochondrial inner membrane that belong to the mitochondrial carrier family (MCF) of proteins. The MCF has many members which act as transporters and display an array of distinct substrate specificities, expression patterns and subcellular localization (Palmieri et al., 2011; Haferkamp & Schmitz-Esser, 2012). After the initial reports of UCP1 (Maia et al., 1998) and UCP2 (Watanabe et al., 1999) as plant homologues of mammalian UCP1, a total of six Arabidopsis isogenes were named *PLANT UNCOUPLING MITOCHONDRIAL PROTEIN* (*PUMP*) *1–6* (Borecký et al., 2006). However, PUMP4–6 exhibit biochemical properties typical of the phylogenetically closely related Mitochondrial Dicarboxylate Carrier (DIC) proteins known from yeast and animals (Palmieri et al., 2008). Accordingly, PUMPs were re-grouped into plant UCP1–UCP3 and plant DIC1–DIC3 (Figure S1) (Palmieri et al., 2008). UCP1 and UCP2 are highly similar in sequence and share 72% amino acid identity, whereas UCP3 only shares 35% and 37% identity with UCP1 and UCP2, respectively (Figure S2A) (Monné et al., 2018).

The biological and molecular functions of plant UCPs have remained a matter of debate since their discovery (Barreto et al., 2020). Measurements in purified coupled mitochondria and functional reconstitution confirmed *bona fide* proton gradient uncoupling activity of Arabidopsis and potato UCP1 (Ježek et al., 1996; Jarmuszkiewicz et al., 1998; Considine et al., 2003; Smith et al., 2004; Borecký et al., 2006; Sweetlove et al., 2006; Ito-Inaba et al., 2008). Similar to mammalian UCP1, the proton conductance of plant UCP1 was activated by superoxide and aldehyde products of lipid peroxidation and inhibited by purine nucleotides (Considine et al., 2003; Smith et al., 2004; Sweetlove et al., 2006). Despite several independent lines of evidence for proton gradient uncoupling activity of plant UCP1, the existence of such an activity was recently challenged based on the *in vitro* reconstitution of UCP1 and UCP2 in phospholipid vesicles (Monné et al., 2018). Electroneutral transport function for dicarboxylates, amino acids, phosphate, sulfate, and thiosulfate was suggested instead. A function in organic acid transport is plausible considering the close relation to DIC proteins (Palmieri et al., 2011; Haferkamp & Schmitz-Esser, 2012; Lee et al., 2021) and given that the uncoupling mechanism of UCPs likely relies on fatty acid shuttling (Vercesi et al., 2006; Berardi & Chou, 2014; Crichton et al., 2017). In this context, it was proposed that both UCP1 and UCP2 are mitochondrial inner membrane transporters important for metabolite exchange between mitochondria and cytosol, possibly to support photorespiratory fluxes. A role for UCP1 in photorespiration and photosynthetic performance is consistent with previous observations in Arabidopsis, tomato and tobacco (Sweetlove et al., 2006; Begcy et al., 2011; Chen et al., 2013; Liu et al., 2015; Barreto et al., 2017; Barreto et al., 2022). Recently, a distinct role for UCP1 in reductive ER stress alleviation was found in roots (Fuchs et al., 2022). In addition, double knockdown lines for the Arabidopsis UCPs, i.e. *ucp1 ucp2, ucp1 ucp3* and *ucp2 ucp3* showed marked growth delay under normal growth conditions while increased sensitivity to salt and osmotic challenge was observed for *ucp1 ucp3* only (Lima et al., 2022). The majority of the studies have linked the plant phenotypes to proton gradient uncoupling activity of UCP1 rather than electroneutral exchange of metabolites and have provided a coherent picture. In this regard, ruling out proton gradient uncoupling activity and instead proposing organic acid transport as the sole function as suggested by Monné et al. (2018) deserves further experimental investigation under conditions closer to the *in vivo* situation.

A prerequisite for a more detailed analysis of UCP function will be clarity about the cellular and physiological context under which the proteins operate. While the localization of plant UCP1 in the inner mitochondrial membrane (IMM) has been consistently confirmed, UCP2 subcellular localization has remained less well defined. Even though the assumption of mitochondrial localization for UCP2 appears parsimonious based on the high degree of amino acid identity when compared to UCP1 as well as similar predicted protein structures (Figure S2A, B), UCP2 is predicted not to reside in the mitochondria (Møller et al., 2020) and rather in the Golgi or endomembrane system by most *in silico* algorithms as summarized by SUBA or Aramemnon (Schwacke et al., 2003; Hooper et al., 2017). Further, UCP2 has not been detected in any published proteomes from purified mitochondrial preparations. Instead, it was detected in a study of the Golgi protein inventory (Parsons et al., 2012) or as a protein proposed to be in the plasma membrane by studies dissecting the Golgi and plasma membrane proteomes (Nikolovski et al., 2012). More recently, a UCP2-GFP construct was found to co-localize with a mitochondrial matrix marker after transient expression under the control of an Arabidopsis *Ubiquitin10* promoter in *Nicotiana benthamiana* leaves followed by protoplast preparation and confocal microscopy of the protoplasts (Monné et al., 2018). Together, these results highlight a discrepancy in the localization of UCP2 and, in turn, in our current understanding of the function of this protein. The importance of assembling multiple lines of evidence in defining the subcellular location of a protein has been a recuring theme in development of our understanding of the complex differences between targeting, accumulation and function in defining subcellular localisation in plants (Millar et al., 2009). Here, we consider several independent lines of evidence including *in silico* analysis and fluorescent protein tagging in stable transgenic Arabidopsis lines to define subcellular localization of UCP2.

A first reason for why UCP2 has not been detected in any published plant mitochondrial proteome may be its low abundance. *UCP1* is the most highly expressed UCP isoform in Arabidopsis based on mRNA abundance (Figure S3A, D) and its protein product is also particularly abundant in the IMM (about 8445 copies per mitochondrion, making up about 1% of IMM surface) (Fuchs et al., 2019). However, *UCP2* and *UCP3* transcript levels are similar, and UCP3 was found in mitochondrial proteomes (about 150 copies per mitochondrion) (Fuchs et al., 2019) along with mitochondrial membrane transport proteins of lower transcript levels than those of *UCP2* and *UCP3* (Figure S3B, E). Putative mitochondrial transporters that were not yet detected in mitochondrial proteomes show typically lower mRNA expression values than *UCP2* (Figure S3C, F). A second reason for the absence of UCP2 in mitochondrial proteomics studies may be because the protein is only present in specific tissues. Mitochondrial proteomes with high coverage were obtained from cell suspensions (Klodmann et al., 2011), whole seedlings (Wagner et al., 2015a), or rosette leaves (Senkler et al., 2017), and have hence covered several but not all possible conditions and tissue types with an appropriate enrichment. While this opens the possibility that UCP2 is present in the mitochondrial proteome of tissues that have not been amenable to specific isolation of mitochondria, the development of cell-type specific mitochondrial isolations through genetic affinity tags (Boussardon et al., 2020; Kuhnert et al., 2020; Lang et al., 2020; Niehaus et al., 2020) provides a tool to address this limitation in the future. However, UCP2’s expression across green tissues and its detection in at least some of the published mitochondrial proteomes would be expected if UCP2 was indeed associated with photorespiratory function as proposed by Monné et al., (2018). The argument that UCP2 peptides may be difficult to detect by the available mass spectrometry approaches can be dismissed considering the high degree of primary sequence identity to UCP1 and the ready detection of UCP2 peptides in two preparations enriched for Golgi and plasma membrane (Nikolovski et al., 2012; Parsons et al., 2012.). While the proteomic evidence is consistent, proving absence is notoriously difficult to achieve. Hence, we have sought orthogonal sources of evidence to shed light on UCP2 localization.

*UCP* transcript expression is relatively stable throughout development, under most stress conditions, and in distinct *Arabidopsis thaliana* accessions (Figure S3A, D) (Clifton et al., 2005; Nogueira et al., 2011). There are exceptions, however, such as seed germination when expression of *UCP1* is increased, similar to many genes encoding mitochondrial proteins, to support mitochondrial biogenesis. In contrast, *UCP2* is not increased (Barreto et al., 2022). To exploit expression patterns systematically and to shed light on subcellular localization, we assessed the co-expression landscapes of *UCP1-3* based on the compilations of RNAseq datasets available in AttedII, Genevestigator and in distinct ecotypes of *Arabidopsis thaliana* (Hruz et al., 2008; Kawakatsu et al., 2016; Obayashi et al., 2018). *UCP1* was part of co-expression clusters containing genes of confirmed mitochondrial localization, while *UCP2* was part of a cluster of genes annotated as predominantly endomembrane and secretory pathway proteins (Figure 1A; Table S1A-D). This picture was confirmed by considering the subcellular location and functional categories of the top 30 co-expressed sets of genes (Hooper et al., 2017). *UCP1* is co-expressed with multiple components of the mitochondrial electron transport chain and the TCA cycle, indicating its expression is linked to that of other proteins involved in mitochondrial energy metabolism (21 of top 30 co-expressed genes are mitochondrial proteins, SUBA4) (Figure 1A, Table S1A, B). In contrast, the *UCP2* co-expression landscape is dominated by nucleotide/sugar transporter protein family gene members, contains no genes associated with mitochondrial energy metabolism, but instead endomembrane and Golgi proteins (16 of the top 30 are confirmed endomembrane proteins, 10 of them confirmed Golgi proteins, only one protein of confirmed mitochondrial localization, SUBA4) (Figure 1A, Table S1C, D). Further, gene ontology enrichment of the transcripts co-expressed with *UCP1* shows an enrichment in processes related to the mitochondrial electron transport chain and TCA cycle (Figure S4A, Table S1E) while for *UCP2* transcripts associated with the Golgi are enriched (Figure S4B, Table S1F). In agreement, genes that are co-expressed with *UCP2* in datasets available through Genevestigator (Table S1G) or in natural accessions (Table S1H) are also enriched with Golgi-targeted genes (Table S1I, J).

**Figure 1.**
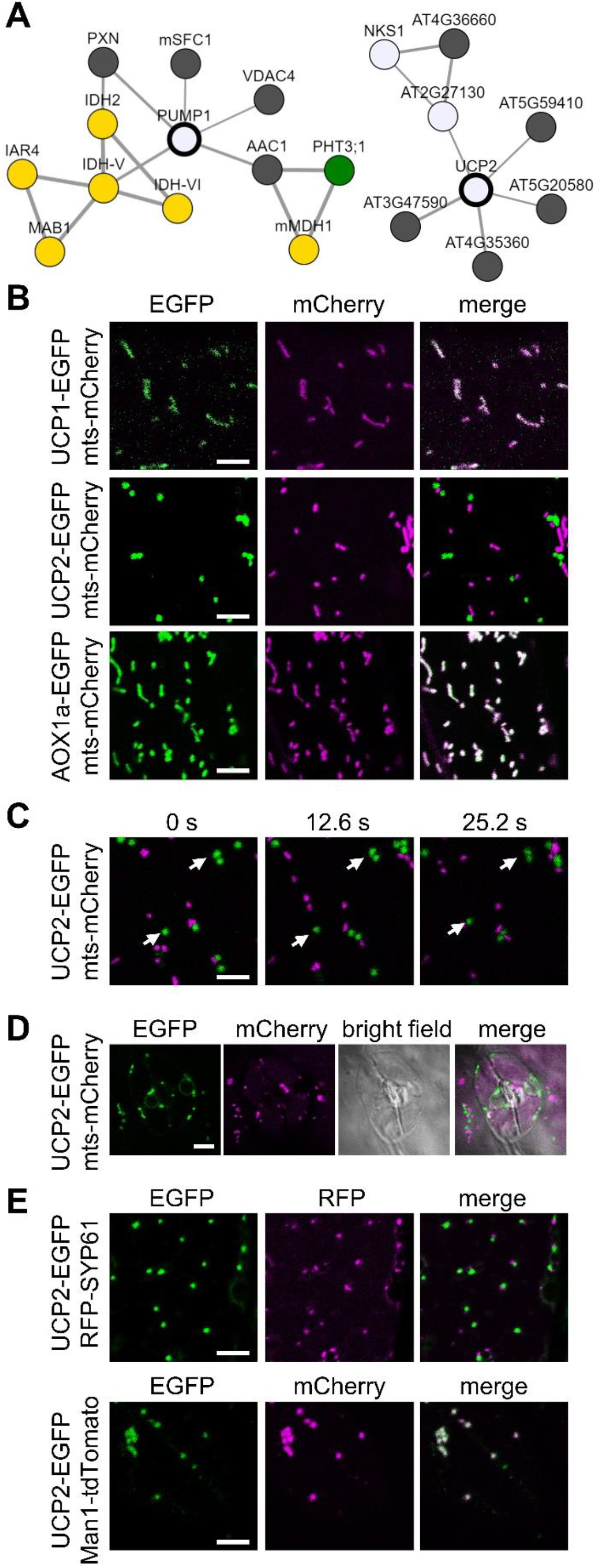
Co-expression landscapes of Arabidopsis UCP1 and UCP2, and CLSM images of seedlings stably co-expressing candidate proteins fused to green fluorescent protein and red fluorescent organellar marker proteins. **(A)** UCP1/PUMP1 (AT3G54110) and UCP2 (AT5G58970) were analysed using co-expression viewer and network drawer tools of AttedII was visualised in the app using Cytoscape. Yellow circles: proteins targeted to the mitochondria; Dark grey circles: proteins targeted to the endomembrane system; Light grey circles: proteins targeted to the secretory pathway; Green circles: proteins targeted to the chloroplasts. Confocal laser scanning microscope (CLSM) images recorded from 3–5 day-old Arabidopsis seedlings stably expressing red fluorescent proteins (mCherry, tdTomato or Red Fluorescent Protein) targeted to different organelles and different candidate proteins fused to enhanced Green Fluorescent Protein (EGFP) under the control of the CaMV35S promoter. EGFP fluorescence: green, mCherry/RFP fluorescence: magenta. (**B**) CLSM images recorded from hypocotyl cells expressing mitochondrial matrix-targeted (mts) mCherry and (top) UCP1-EGFP, (middle) UCP2-EGFP or (bottom) AOX1a-EGFP. (**C**) CLSM time course of a hypocotyl cells expressing mts-mCherry and UCP2-EGFP. Individual images represent merged channels of EGFP and mCherry. White arrows indicate static EGFP-labelled structures relative to dynamic mCherry-labelled mitochondria. The complete time course is shown in Video S1. (**D**) CLSM images of two guard cells at the abaxial epidermis of cotyledons expressing mts-mCherry and UCP2-EGFP. (**E**) CLSM images recorded from hypocotyl cells expressing UCP2-EGFP and targeting sequence of soybean Mannosidase1 fused to N-terminus of tdTomato (Man1-tdTomato; Golgi marker) or Arabidopsis syntaxin protein 61 fused to the C-terminus of RFP (RFP-SYP61; trans-Golgi network/Early Endosome marker). Scale bars = 5 μm.

We next investigated the localization of UCP2 using an orthogonal cell biological approach. Enhanced GFP (EGFP) was fused to the C-terminus of the coding sequence of *UCP2* splice form 1 under the control of the Cauliflower mosaic virus 35S (CaMV35S) promoter and transformed into an Arabidopsis line expressing mCherry targeted to the mitochondrial matrix (mts-mCherry) (el Zawily et al., 2014) to obtain a stable dual marker line.

A second splice form of *UCP2* that was predicted *in silico* is identical to splice form 1 except for a shorter C-terminus. Since predicted splice form 2 may in principle show differential subcellular targeting, we asked whether this splice form is actually produced *in vivo*, or more plausibly a prediction artefact. Predicted splice form 2 could not be amplified from seedling cDNA and there was no evidence of this putative splice form in any of the available RNAseq public datasets or as protein based on peptide data. Together, this suggests that splice form 1 is either the dominant form of UCP2 by far while splice form 2 is confined to a cell type that has not been covered by dedicated analyses, or splice form 1 is the sole splice form expressed. Following the principle of parsimony, we focussed our further analysis on splice form 1. Confocal microscopy of hypocotyl cells from seedlings revealed co-localization of UCP1-EGFP and a known mitochondrial marker, Alternative Oxidase 1A-EGFP (AOX1A-EGFP) respectively, with mCherry-labelled mitochondria (Figure 1B). In contrast, EGFP signal from the fusion with UCP2 revealed punctate fluorescent structures of regular, disc-like shape that did not colocalize with mitochondria (Figure 1B). Time lapse analysis of the UCP2-EGFP structures further confirmed distinct movement from the mCherry-labelled mitochondria (Figure 1C, Video S1). While both distinct structures moved in a directed manner, the EGFP-labelled structures were less motile than the mitochondria. The intracellular velocity of mitochondria in seedlings (3 to 6 days after germination) has been estimated to be approximately double than that of Golgi stacks or peroxisomes, which are of similar size to mitochondria (Stefano et al., 2014).

The absence of co-localization with mCherry-labelled mitochondria was not only found in hypocotyl epidermal cells but was consistent in all cell types assessed, including leaf guard cells (Figure 1D). Additional residual EGFP signal in the secretory system as evident from nuclear ring structures was also observed. Similar nuclear envelope-fluorescent structures are regularly observed when overexpressing GFP-fusions of proteins of the secretory system, including Golgi-resident proteins, such as Synaptotagmin 2 fused to GFP (Zhang et al., 2011). To test whether the labelled structures really represented Golgi localization, we next transformed the UCP2-EGFP construct into Arabidopsis marker lines expressing the Golgi marker Man1-tdTomato, consisting of the first 49 amino acids of *Glycine max* α-1,2-MANNOSIDASE (Nelson et al., 2007) or the *trans*-Golgi Network/Early Endosome (TGN/EE) marker RFP-SYP61 (Henderson et al., 2015). Using stable lines for imaging, UCP2-EGFP signal co-localized with Man1-tdTomato signal (Figure 1E), demonstrating that UCP2-EGFP localizes to the Golgi. UCP2-EGFP signal was systematically found in the direct vicinity of RFP-SYP61 labelled structures, further supporting UCP2 localization in the Golgi but not the TGN/EE (Figure 1E).

To test whether Golgi localization of UCP2 is conserved in eudicots beyond Arabidopsis, we devised co-localization experiments using the tomato (*Solanum lycopersicum*) orthologues of Arabidopsis UCP1 and UCP2 (hereafter referred to as SlUCP1 and SlUCP2; Figure S1 and Figure S5). C-terminal EGFP fusions of SlUCP1 and SlUCP2 were transiently expressed in *N. benthamiana* leaves and co-infiltrated with a SlAOX1a-RFP marker construct for mitochondrial localization. SlUCP1-EGFP co-localized with RFP fluorescence, validating mitochondrial localization, while SlUCP2-EGFP showed distinct localization from the RFP signal (Figure S6). The EGFP signal showed a punctuate pattern of compartments similar to that found for *Arabidopsis thaliana* UCP2-EGFP (Figure 1B-E).

Recent experiments using *ucp2* lines (Monné et al., 2018; Arcuri et al., 2021; Lima et al., 2022) deserve careful re-interpretation and future reverse genetic strategies to understand UCP function in mitochondria should employ *ucp1* and *ucp3* lines and their crosses. It has been shown that some members of the MCF protein family are not localized in mitochondria, for instance specific MCF family members are found in plastids (Bouvier et al., 2006; Bahaji et al., 2011; Gigolashvili et al., 2012), peroxisomes (Fukao et al., 2001; Arai et al., 2008; Linka et al., 2008), the plasma membrane (Rieder & Neuhaus, 2011) and in the endomembrane system (Leroch et al., 2008; Chu et al., 2017). The remarkably high amino acid identity between UCP1 and UCP2 but differing subcellular localization may imply that a proportion of the difference serves as targeting information, which make UCP1 and UCP2 a valuable model system to reveal novel subcellular targeting principles in future studies. The similarity between UCP1 and UCP2 amino acid sequences and predicted protein structures (Figure S2 and Figure S5) from Arabidopsis and tomato suggest a similar mode of UCP-mediated transmembrane transport in both the IMM and the Golgi membranes, despite physiologically different circumstances (e.g. in electrical potential or metabolite concentrations at both surfaces). Yet, both membranes share pH gradients (approximately 0.5 pH units across the Golgi membrane and 0.8 pH units across the IMM) (Martinière et al., 2013; Shen et al., 2013). Therefore, proton-coupled transport through UCP2 in the Golgi membrane is a possibility.

We conclude that reverse genetic studies should consider UCP1 and UCP2 separately because the cumulative evidence on the subcellular targeting points to Golgi targeting of UCP2. While taking together the independent lines of evidence provides robustness and coherence, each individual line of evidence is less than definitive when considered in isolation. Co-expression data provide indirect and correlative evidence on targeting, proteomic studies are incomplete in coverage and reflect the characteristics of the prepared subcellular fractions, and fluorescent protein tagging can lead to mistargeting as a consequence of the tagging or altered promoter activity and transcript dynamics. Even though the current evidence supports Golgi targeting of UCP2, it cannot definitively rule out that UCP2 might localize to mitochondria under specific conditions. That being said, there is currently no evident concept for conditional targeting of transmembrane proteins between the mitochondria and Golgi. Future investigations of the endomembrane system should consider UCP2 function in the Golgi as an interesting player in the context of the activity of other Golgi resident transport proteins to set pH and ion homeostasis in the Golgi lumen (Scholl et al., 2021; McKay et al., 2022).

## METHODS

### Plant material

*Arabidopsis thaliana* co-expressing EGFP and mCherry/tdTomato/RFP fusion constructs were generated by Agrobacterium-mediated transformation via floral dip as previously described (Steinbeck et al., 2020). Arabidopsis stably transformed to express mts-mCherry (el Zawily et al., 2014), a kind gift from David Logan, were transformed with UCP1-EGFP, UCP2.1-EGFP or AOX1a-EGFP expression constructs. Arabidopsis lines stably transformed to express Man1-tdTomato and RFP-SYP61 were transformed with the UCP2-EGFP expression construct. The Man1-tdTomato line carries the first 49 amino acids of Soybean Mannosidase 1 fused with tdTomato (Nebenführ et al., 1999). Lines co-expressing EGFP and mCherry/tdTomato/RFP were selected at the stereomicroscope employing a GFP filter (excitation: 470 ± 20 nm, emission: 525 ± 25 nm) and a DsRed filter (excitation: 545 ± 15 nm, emission: 620 ± 30 nm), respectively.

### Cloning

Gene transcripts encoding Arabidopsis (UCP1, UCP2.1 and AOX1a) and Tomato (SlUCP1, SlUCP2 and SlAOX1a) proteins were enriched using primers binding in UTRs to serve as template for second PCR adding *att*B Gateway recombination sites. Attempts of amplifying the predicted splice variant UCP2.2 from Arabidopsis did not result in a PCR product. Amplified DNA fragments were purified and BP-recombined into the pDONR207 Gateway donor vector (Invitrogen). Accuracy of the inserted sequence and assembly was confirmed by DNA sequencing. Verified recombined pDONR207 vectors containing the Arabidopsis sequences were LR-recombined with modified pSS01 destination vectors. The roGFP2 (in frame behind the Gateway cassette) of the pSS01 vector (Brach et al., 2009) was replaced with EGFP using AvrII and PacI cut/ligation. The pUBC-EGFP-Dest vector (Blatt & Grefen, 2014) was used as DNA template for PCR amplification of EGFP. After recombination of the pSS01 destination with the recombined pDONR207 vectors, correct assembly was confirmed by DNA sequencing. Verified recombined pDONR207 vectors containing SlUCP1, SlUCP2 and SlAOX1a were LR-recombined with PGWB405 (EGFP) and pGWB454 (RFP) (Nakagawa et al., 2007). Primers used for cloning procedures, sequencing and vector backbone modifications are listed in Table S2.

### Confocal laser scanning microscopy

Arabidopsis imaging with confocal laser scanning microscopy (CLSM) was performed as described previously (Wagner et al., 2015b) using either a Zeiss LSM780 or LSM980 confocal microscope and a ×63 lens (Plan-Apochromat, 1.40 N.A., oil immersion). EGFP was excited at 488 nm and fluorescence collected at 495–545 nm. mCherry/tdTomato/RFP was excited at 543 nm and fluorescence collected at 570–624 nm. An Olympus FV1000 confocal laser microscope with a ×60 lens (U-PlanSApo, 1.2 N.A., water immersion) was used for the observation of the subcellular localization of tomato proteins in leaf sections of *Nicotiana benthamiana* plants. EGFP was excited at 488 nm and fluorescence collected at 490-550 nm. RFP was excited at 559 nm and emission was collected at 570–650 nm.

### Phylogenetic analysis

Protein sequences of representative species were retrieved from UniProt Standard Protein Blast using Arabidopsis UCP1 (UniProt identifier O81845) as query sequence. After manual curation, sequences were aligned with MUSCLE (Edgar, 2004) in MEGA11 (Stecher et al., 2020) with default parameters. An unrooted tree was calculated using MrBayes with split frequencies <0.01 after 2,000,000 generations with a mixed amino acid model (Huelsenbeck & Ronquist, 2001). Sequences and accessions used for phylogenetic analysis are available in Table S3.

### In silico analysis

Protein alignments shown in Figure S2 and S5 were performed with Multalin (Corpet, 1988.) and images were generated with ESPript 3.0 (Robert & Gouet, 2014). Protein structures were modelled with PyMol (The PyMOL Molecular Graphics System, Version 1.2r3pre, Schrödinger, LLC) and Alphafold (Jumper et al., 2021). Co-expression of UCP1/PUMP1 (AT3G54110), UCP2/PUMP2 (AT5G58970) and UCP3/PUMP3 (AT1G14140) were analysed using co-expression viewer and network drawer tools of AttedII (Obayashi et al., 2018) or Genevestigator (Hruz et al., 2008). Network drawer used ath-m. c9-0 and was visualised in the app using Cytoscape. Sub-cellular localization data for the top 30 co-expressed genes for each UCP gene (as defined by CoExSearch on AttedII) were obtained from SUBA4 (Hooper et al., 2017). Transcriptomic data from a collection of natural accession was obtained from 1,001 Genomes Project (Kawakatsu et al., 2016) and analyzed using Genevestigator (Hruz et al., 2008).

## ACKNOWLEDGEMENTS

This work was supported by the Deutsche Forschungsgemeinschaft (DFG, German Research Foundation) through the Research Training Group GRK 2064 “Water use efficiency and drought stress responses: From Arabidopsis to Barley” (to A.J.M. and M.S.), the infrastructure grant <u>INST 211/903-1 FUGG</u> for a confocal microscope as operated by the Imaging Network of the University of Münster (RI_00497) and the Spanish Ministry for Science and Innovation (MCIN)/ Agencia Estatal de Investigación (AEI) /10.13039/501100011033 project PID2020-120229RA-I00. E.F.-P. was supported by a pre-doctoral fellowship PRE2021-097120 and I.F.-S. by the ‘Ramon y Cajal’ contract RYC2019-028030-I, both funded by MCIN/ AEI /10.13039/501100011033 and by “ESF Investing in your future”. P.B. was supported by a postdoctoral fellowship 200385/2022-4 funded by the Brazilian National Council for Scientific and Technological Development (CNPq).

## Supplementary Information

**Supplementary Figure 1.**
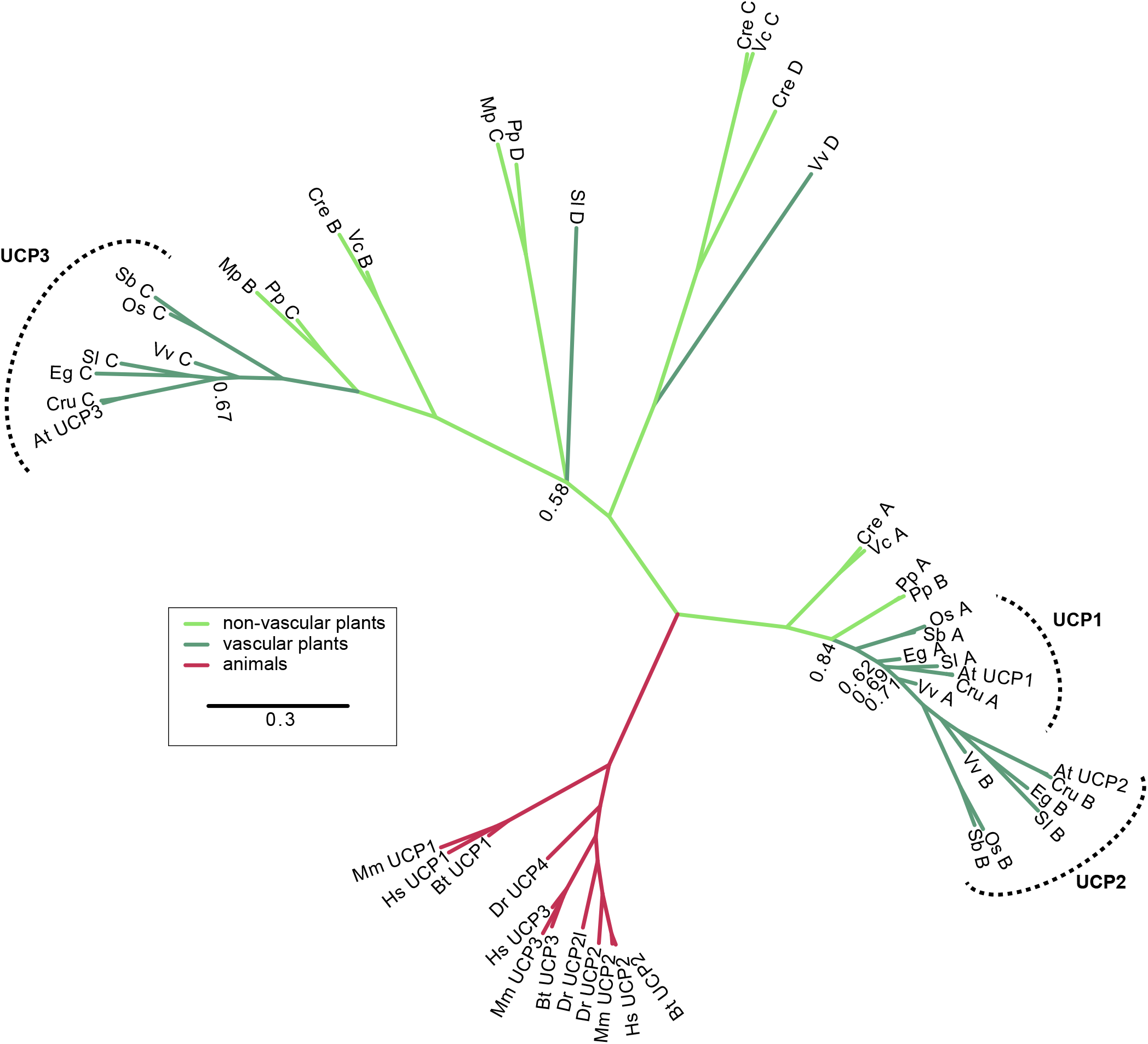
Phylogenetic tree of homologs from representative plant and animal species to *Arabidopsis thaliana* UCP1 protein. Scale bar represents 30 substitutions per 100 amino acids. Numerical values along the tree represent bootstrap values of respective branching (only values < 0.9 shown). The corresponding sequences are shown in Supplementary Table 1. At: *Arabidopsis thaliana*, Bt: *Bos taurus*, Cru: *Capsella rubella*, Cre: *Chlamydomonas reinhardtii*, Dr: *Danio rerio*, Eg: *Erythranthe guttata*, Hs: *Homo sapiens*, Mp: *Marchantia polymorpha*, Mm: *Mus musculus*, Os: *Oryza sativa*, Pp: *Physcomitrium patens*, Sb: *Sorghum bicolor*, Sl: *Solanum lycopersicum*, Vv: *Vitis vinifera*, Vc: *Volvox carteri*.

**Supplementary Figure 2.**
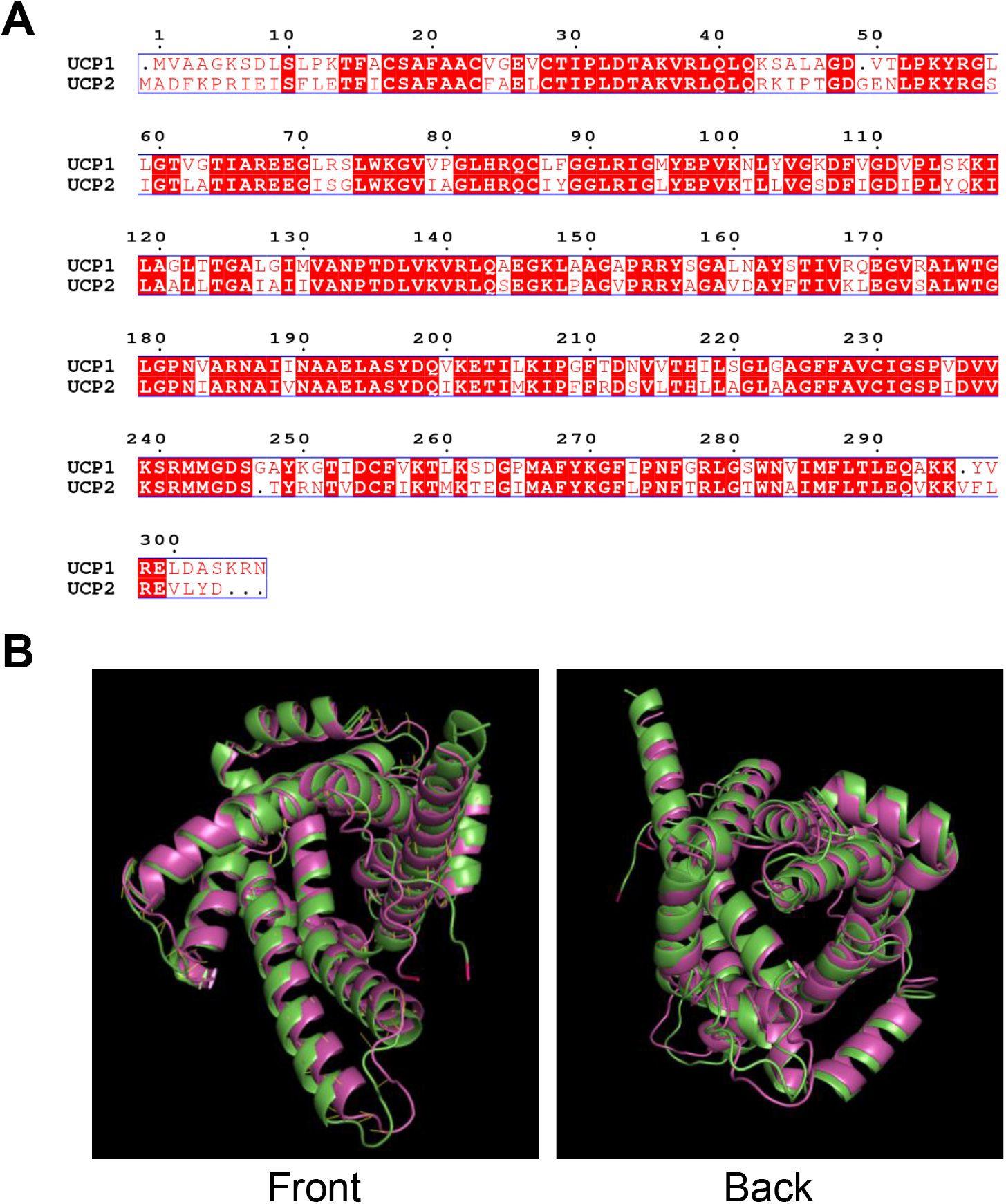
Sequence and structural alignment of Arabidopsis UCP1 and UCP2. **(A)** Multiple sequence alignment of Arabidopsis UCP1 (AT3G54110, NP_190979.1) and UCP2 (AT5G58970, NP_568894.1). **(B)** Front (left) and back (right) views of UCP1 (green) and UCP2 (magenta) protein structural alignment.

**Supplementary Figure 3.**
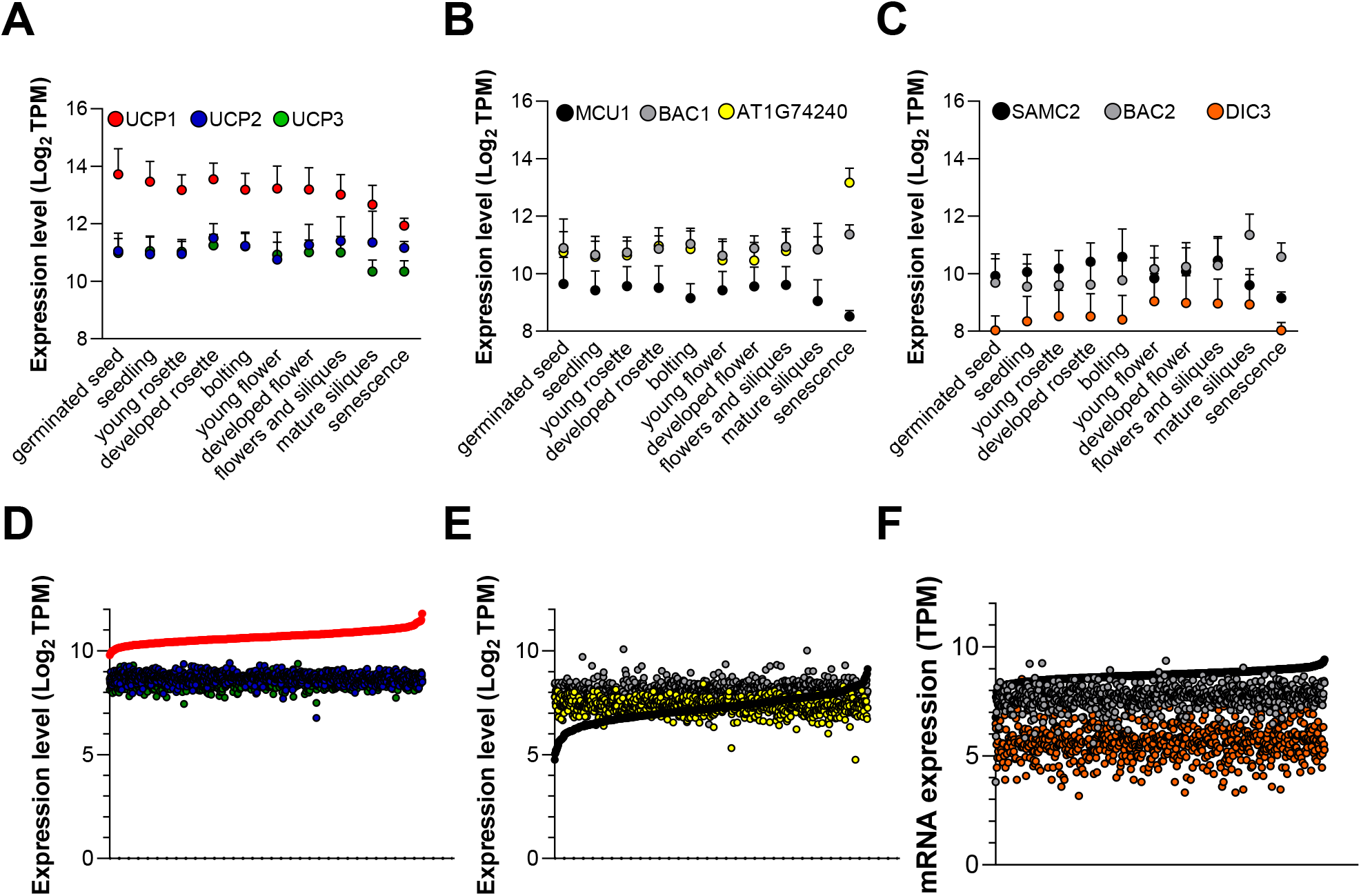
Gene expression of genes annotated as mitochondrial transporters during Arabidopsis development and in a collection of natural accessions. Absolute mRNA abundance values of **(A, D)** UCPs, **(B, E)** low-copy number proteins in the mitochondrial proteome (Fuchs et al., 2019) and **(C, F)** transporters not found in the mitochondrial proteome during *Arabidopsis thaliana* **(A, B, C)** development and in a **(D, E, F)** collection of Arabidopsis accessions. MCU1, AT1G09575; BAC1, AT2G33820; AT1G74240; SAMC2, AT1G34065; BAC2, AT1G79900; DIC3, AT5G09470. **(A, B, C)** Number of samples for each data point is: 515, germinated seed; 2781, seedlings; 830, young rosette; 2219, developed rosette; 358, bolting; 720, young flower; 1003, developed flower; 274, flowers and siliques; 93, mature siliques; 18, senescence. **(D, E, F)** Each data point is the absolute transcript per million (TPM) value from a single accession from a total of 727 accessions plotted along the X axis (Kawakatsu et al., 2016). Data is ordered from the lowest to the highest expression of **(D)** UCP1, **(E)** MCU1 and **(F)** SAMC2.

**Supplementary Figure 4.**
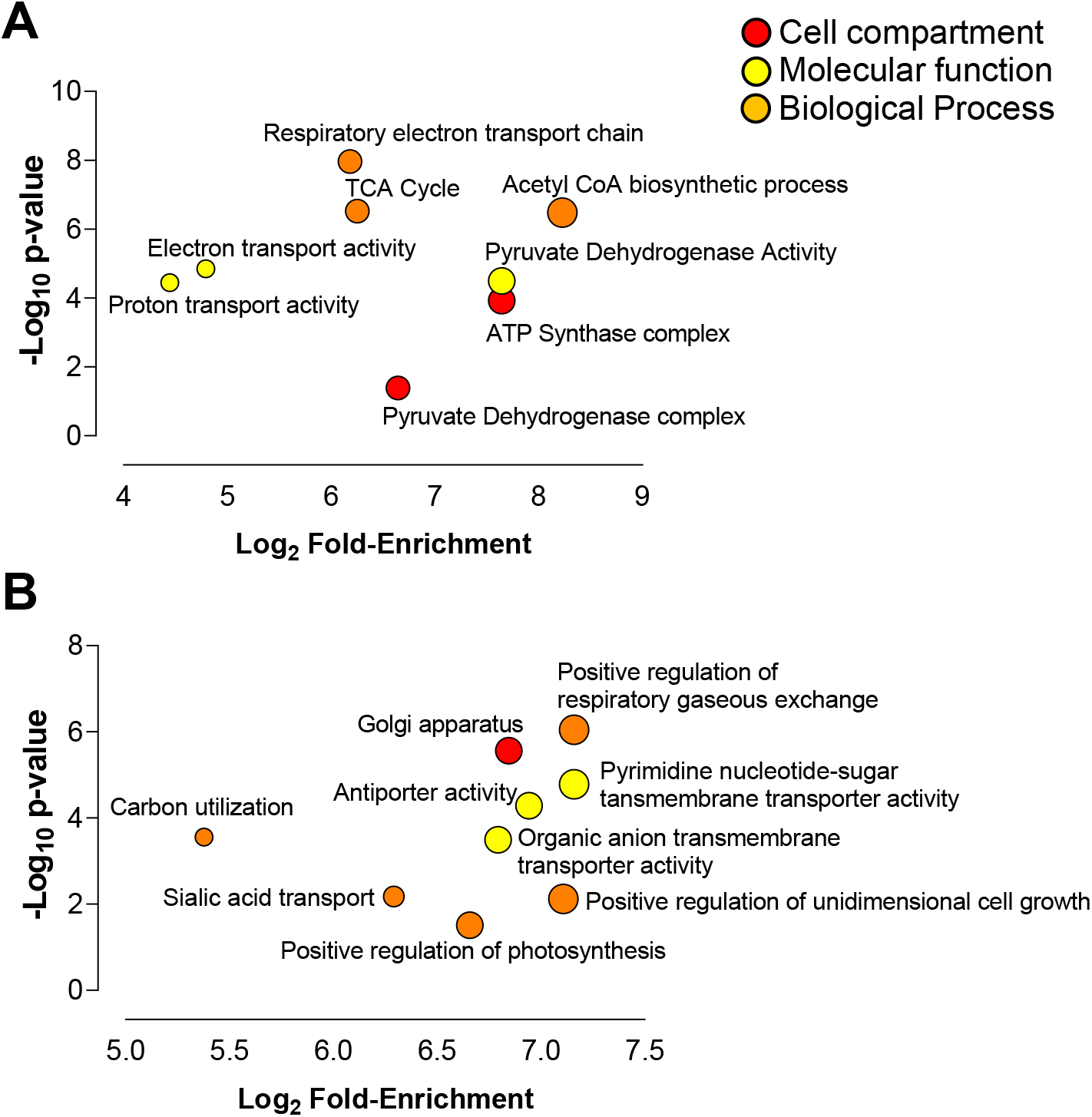
**(A)** The top 30 proteins co-expressed with UCP1 (Table S1A, B) and **(B)** UCP2 (Table S1C, D) were used as queries for a Gene Ontology Term Enrichment analysis against the whole Arabidopsis transcriptome as a reference. Circle areas scale proportionally to Log2 Fold-Enrichment. The full list of GO enriched bins are provided in Table S1E and Table S1F.

**Supplementary Figure 5.**
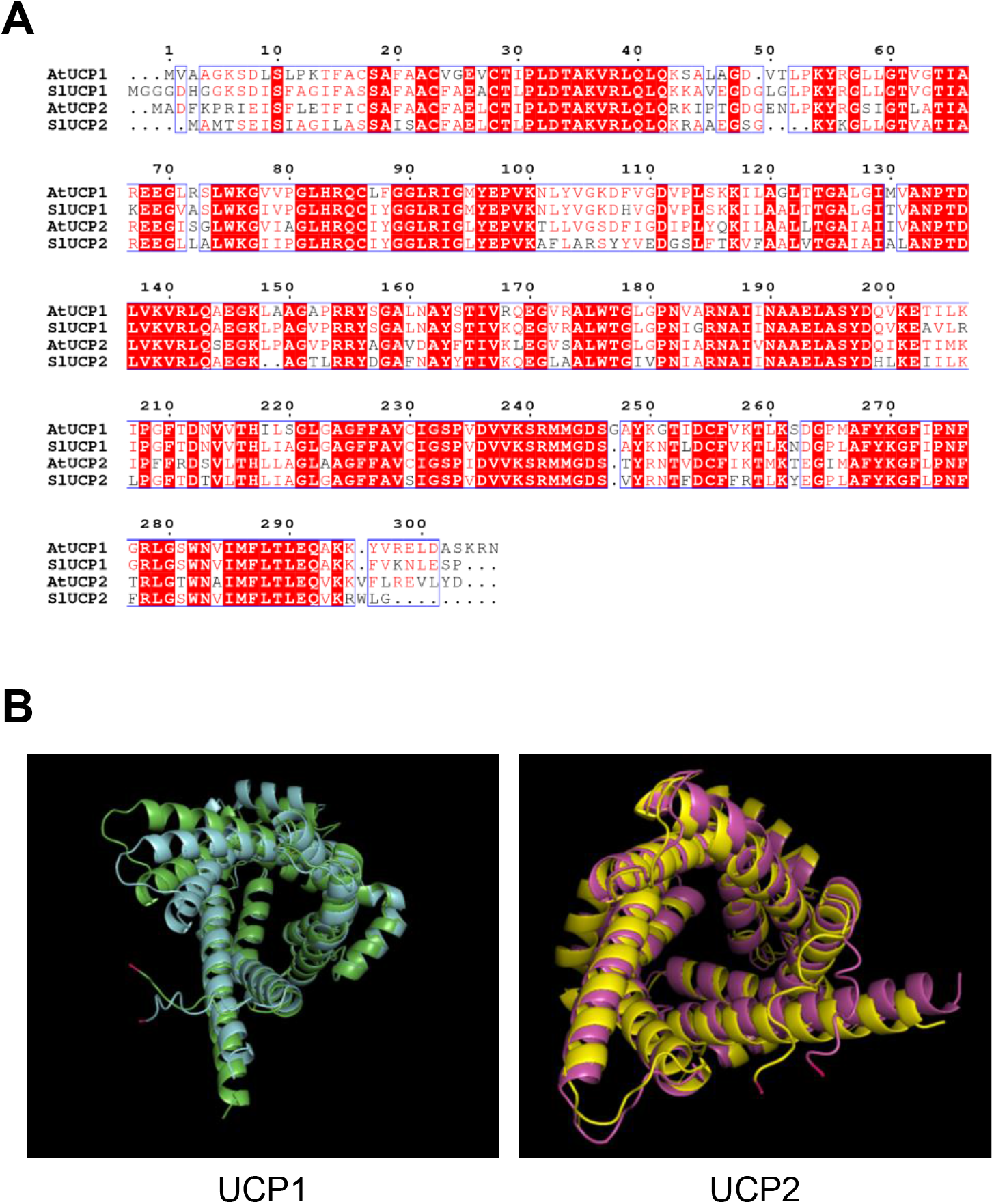
Sequence and structural alignment of UCP1 and UCP2 from Arabidopsis and Tomato. **(A)** Multiple sequence alignment of Arabidopsis (At) UCP1 (AT3G54110, NP_190979.1; green) and UCP2 (AT5G58970, NP_568894.1; magenta) together with Tomato (Sl) UCP1 (Solyc09g011920, NP_001234584.1; blue) and UCP2 (Solyc09g031680, XP_004246961.1; yellow). AtUCP1 has 83.77% and 69.55% identity with SlUCP1 and SlUCP2 respectively. AtUCP2 has 70% and 71.05% identity with SlUCP1 and SlUCP2 respectively. (b) Alignments of the predicted structures for At and Sl UCP1 (left) and UCP2 (right).

**Supplementary Figure 6.**
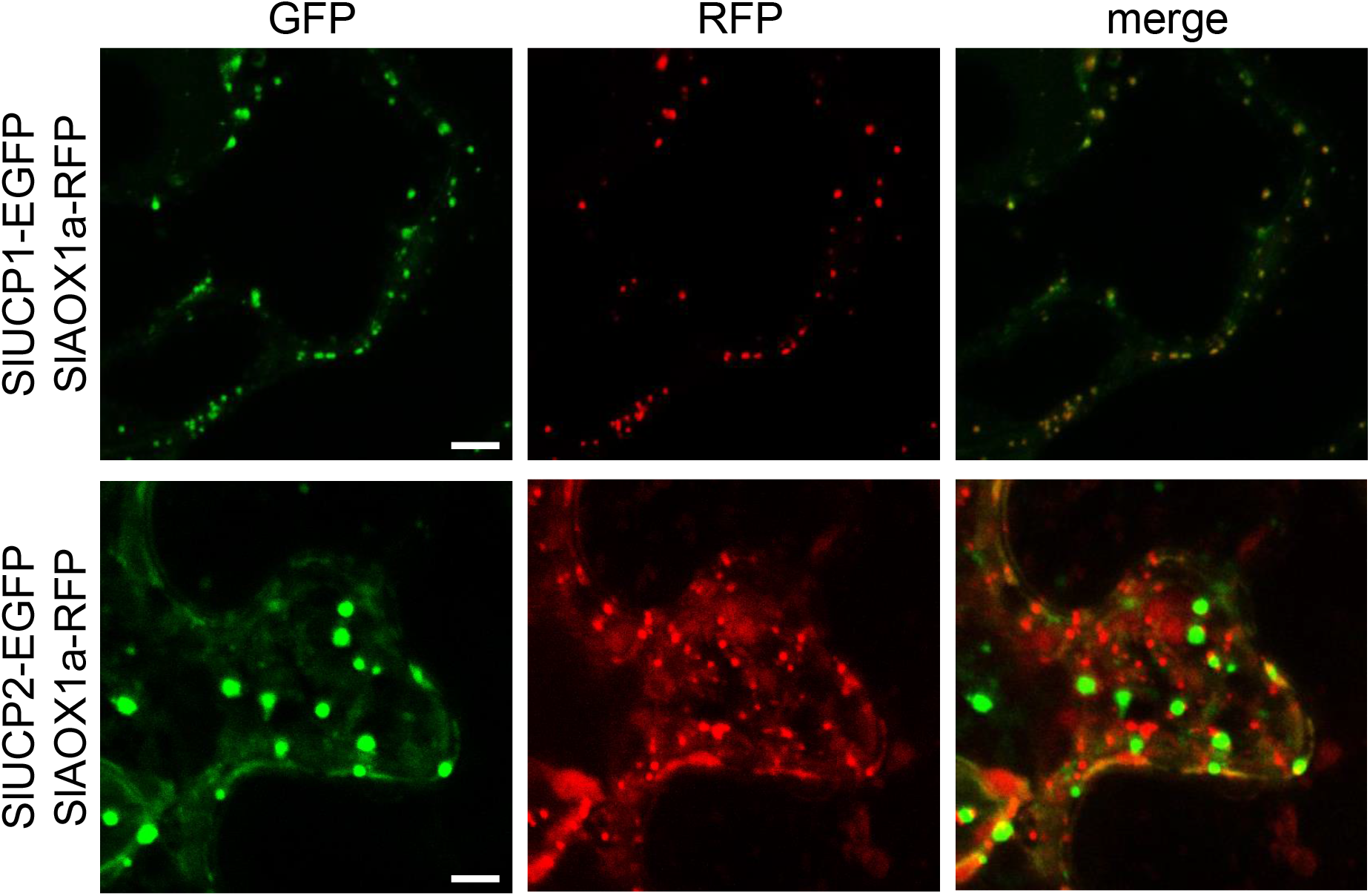
Representative CLSM images of *Nicotiana benthamiana* leaves infiltrated with SlUCP1, SlUCP2 and SlAOX1a fused to fluorescent proteins. Confocal laser scanning microscope images were recorded from leaves 3 days after co-infiltration. Leaves were co-infiltrated with SlUCP1 or SlUCP2 fused to Enhanced Green Fluorescent Protein (EGFP) together with SlAOX1a fused to Red Fluorescent Protein (RFP). Both constructs are under the control of cauliflower mosaic virus 35S promoter. EGFP fluorescence: green, RFP fluorescence: red. Scale bars = 5 μm.

**Supplementary Table 1**. *In silico* analysis of UCP1, UCP2 and UCP3 co-expression landscapes. **(A)** Top 30 Genes co-expressed with UCP1 using AttedII, **(B)** Subcellular prediction for genes listed in Table S1B using SUBA4, **(C)** Top 30 Genes co-expressed with UCP2 using Atted II, **(D)** Subcellular prediction for genes listed in Table S1C using SUBA4, **(E)** TAIR GO Enrichment for genes co-expressed with UCP1 (listed on Table S1A, B), **(F)** TAIR GO Enrichment for genes co-expressed with UCP2 (listed on Table S1C, D), **(G)** Top 30 Genes co-expressed with UCP2 using Genevestigator, **(H)** Genes co-expressed with UCP2 (score ≥ 0.7) across Arabidopsis natural accessions, **(I)** TAIR GO Enrichment for genes co-expressed with UCP2 (Listed on Table S1G) and **(J)** TAIR GO Enrichment for genes co-expressed with UCP2 (Listed on Table S1H).

**Supplementary Table 2**. List of oligonucleotides used for cloning, sequencing and vector backbone modifications.

**Supplementary Table 3**. Sequences and accessions used for phylogenetic analysis.

**Supplemental Video 1**. Time lapse analysis of the UCP2-EGFP.

