## Supplemental Figures for "PLANT UNCOUPLING MITOCHONDRIAL PROTEIN 2 localizes to the Golgi"

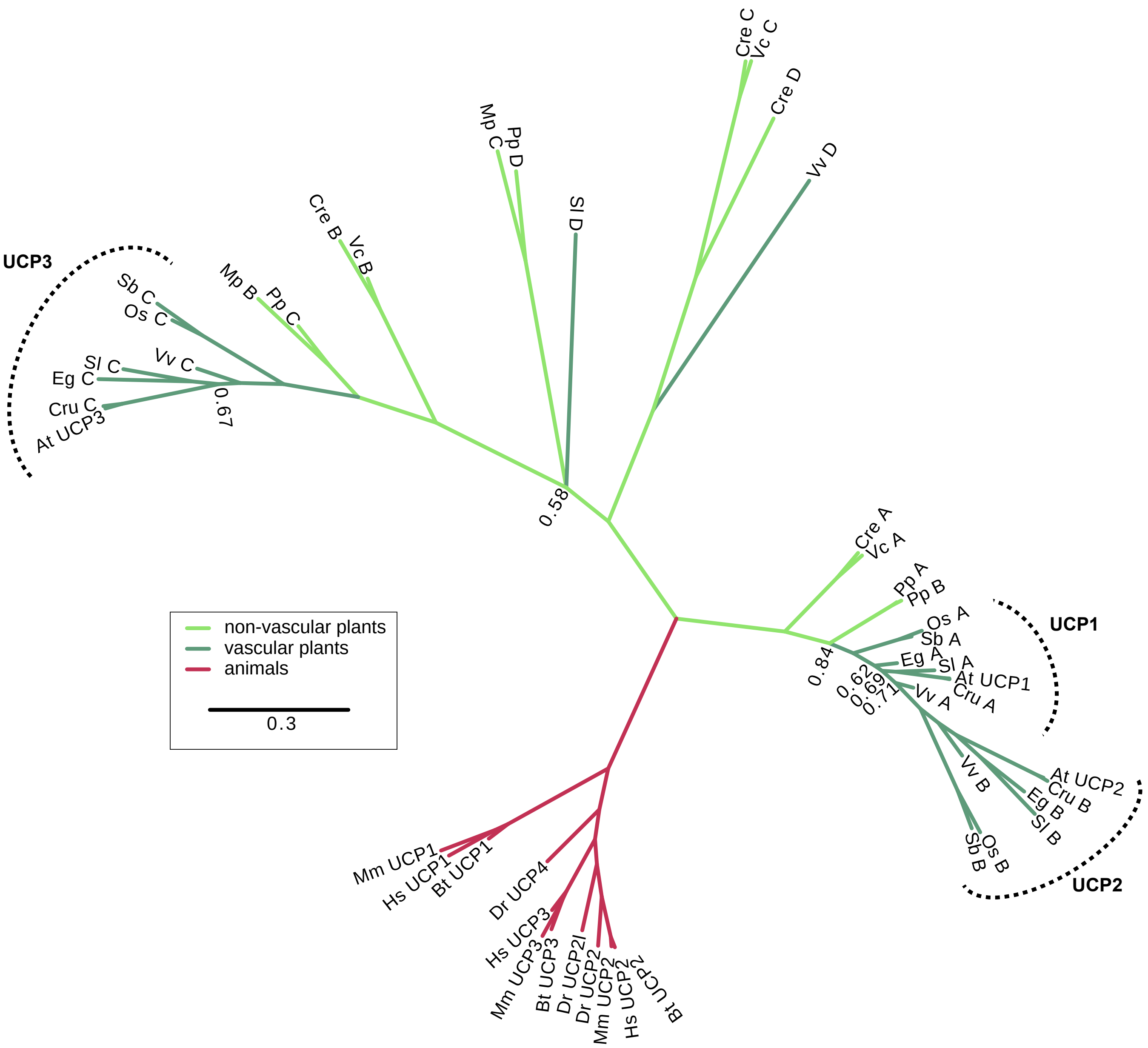


**Supplementary Figure 1.** Phylogenetic tree of homologs from representative plant and animal species to *Arabidopsis thaliana* UCP1 protein. Scale bar represents 30 substitutions per 100 amino acids. Numerical values along the tree represent bootstrap values of respective branching (only values < 0.9 shown). The corresponding sequences are shown in Supplementary Table 1. At: *Arabidopsis thaliana*, Bt: *Bos taurus*, Cru: *Capsella rubella*, Cre: *Chlamydomonas reinhardtii*, Dr: *Danio rerio*, Eg: *Erythranthe guttata*, Hs: *Homo sapiens*, Mp: *Marchantia polymorpha*, Mm: *Mus musculus*, Os: *Oryza sativa*, Pp: *Physcomitrium patens*, Sb: *Sorghum bicolor*, Sl: *Solanum lycopersicum*, Vv: *Vitis vinifera*, Vc: *Volvox carteri*.

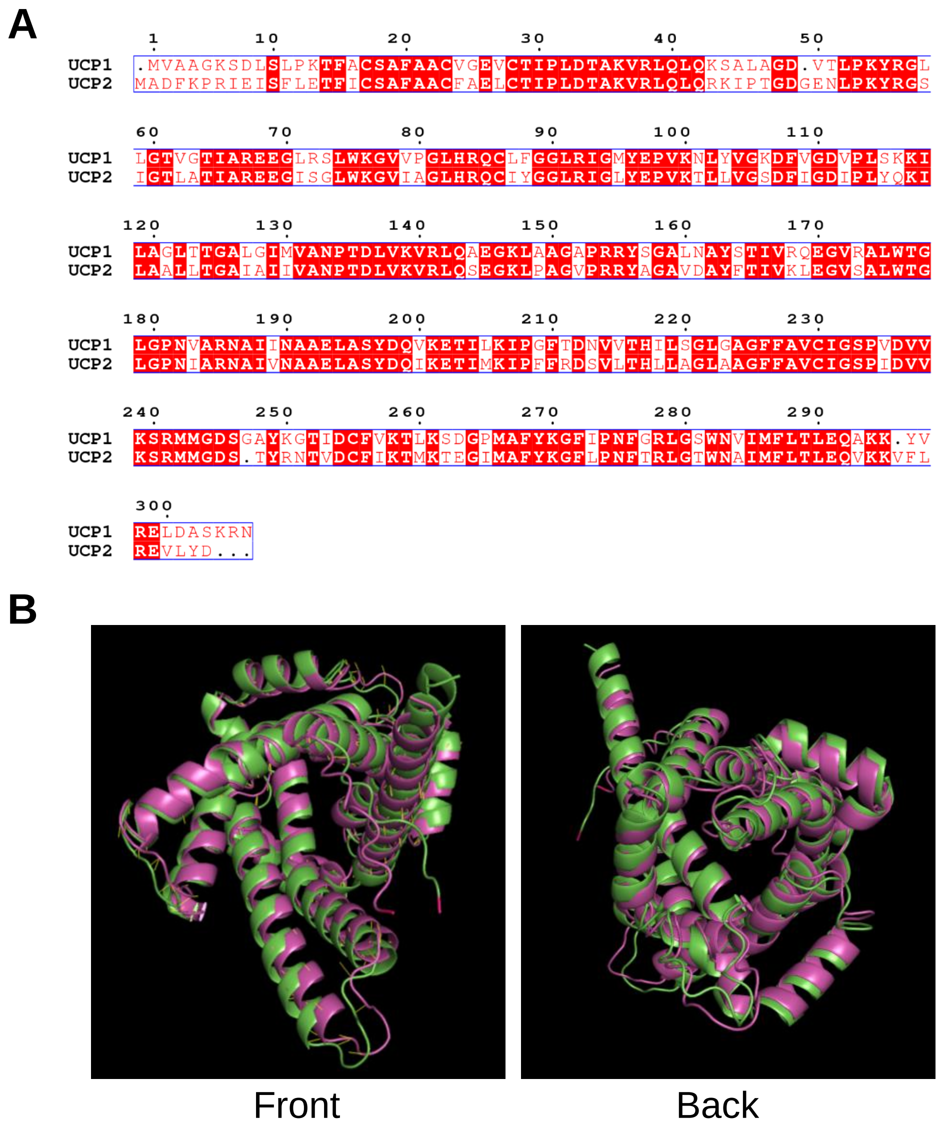


**Supplementary Figure 2.** Sequence and structural alignment of Arabidopsis UCP1 and UCP2. **(A)** Multiple sequence alignment of Arabidopsis UCP1 (AT3G54110, NP_190979.1) and UCP2 (AT5G58970, NP_568894.1). **(B)** Front (left) and back (right) views of UCP1 (green) and UCP2 (magenta) protein structural alignment.

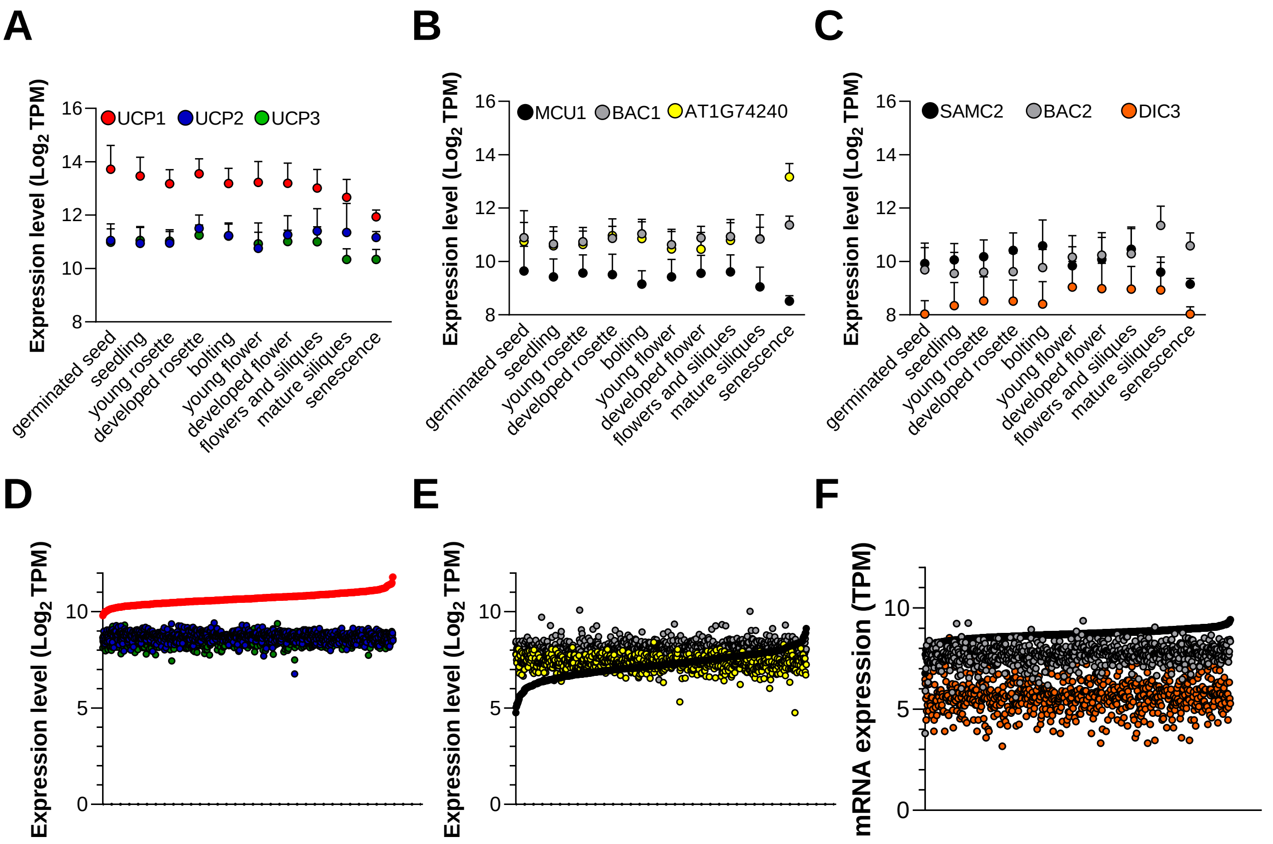


**Supplementary Figure 3.** Gene expression of genes annotated as mitochondrial transporters during Arabidopsis development and in a collection of natural accessions. Absolute mRNA abundance values of **(A, D)** UCPs, **(B, E)** low-copy number proteins in the mitochondrial proteome (Fuchs et al., 2019) and **(C, F)** transporters not found in the mitochondrial proteome during *Arabidopsis thaliana* **(A, B, C)** development and in a **(D, E, F)** collection of Arabidopsis accessions. MCU1, AT1G09575; BAC1, AT2G33820; AT1G74240; SAMC2, AT1G34065; BAC2, AT1G79900; DIC3, AT5G09470. **(A, B, C)** Number of samples for each data point is: 515, germinated seed; 2781, seedlings; 830, young rosette; 2219, developed rosette; 358, bolting; 720, young flower; 1003, developed flower; 274, flowers and siliques; 93, mature siliques; 18, senescence. **(D, E, F)** Each data point is the absolute transcript per million (TPM) value from a single accession from a total of 727 accessions plotted along the X axis (Kawakatsu et al., 2016). Data is ordered from the lowest to the highest expression of **(D)** UCP1, **(E)** MCU1 and **(F)** SAMC2.

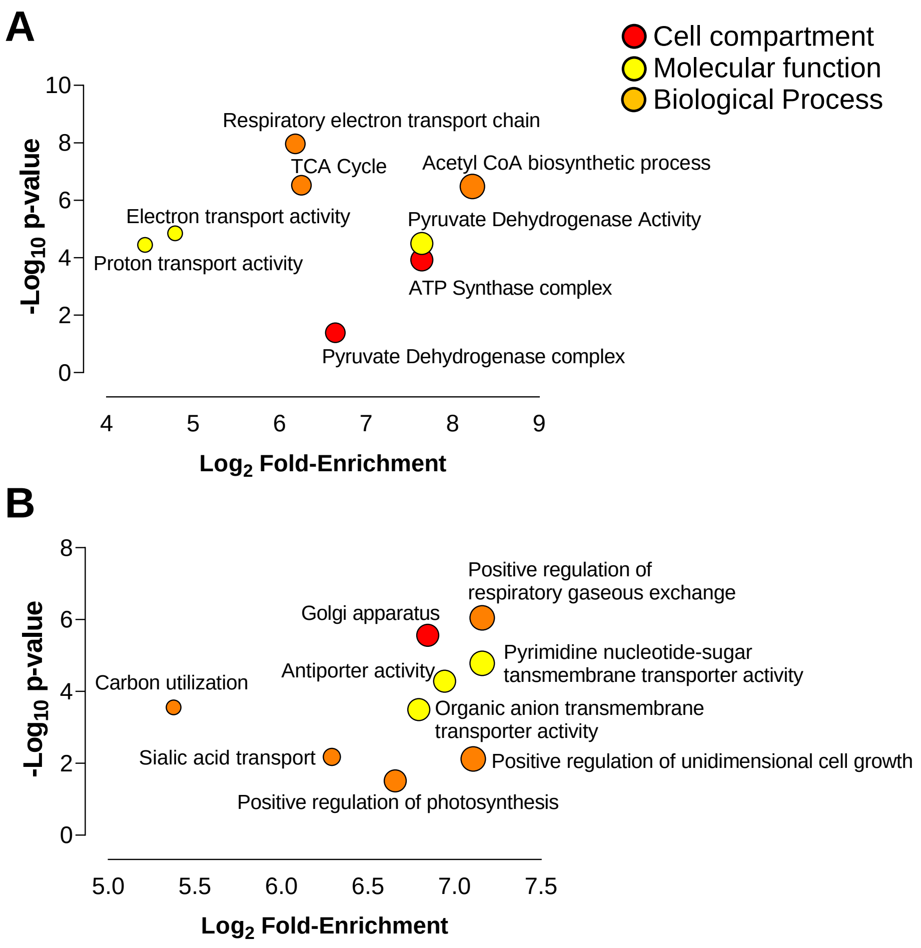


**Supplementary Figure 4. (A)** The top 30 proteins co-expressed with UCP1 (Table S1A, B) and **(B)** UCP2 (Table S1C, D) were used as queries for a Gene Ontology Term Enrichment analysis against the whole Arabidopsis transcriptome as a reference. Circle areas scale proportionally to Log2 Fold-Enrichment. The full list of GO enriched bins are provided in Table S1E and Table S1F.

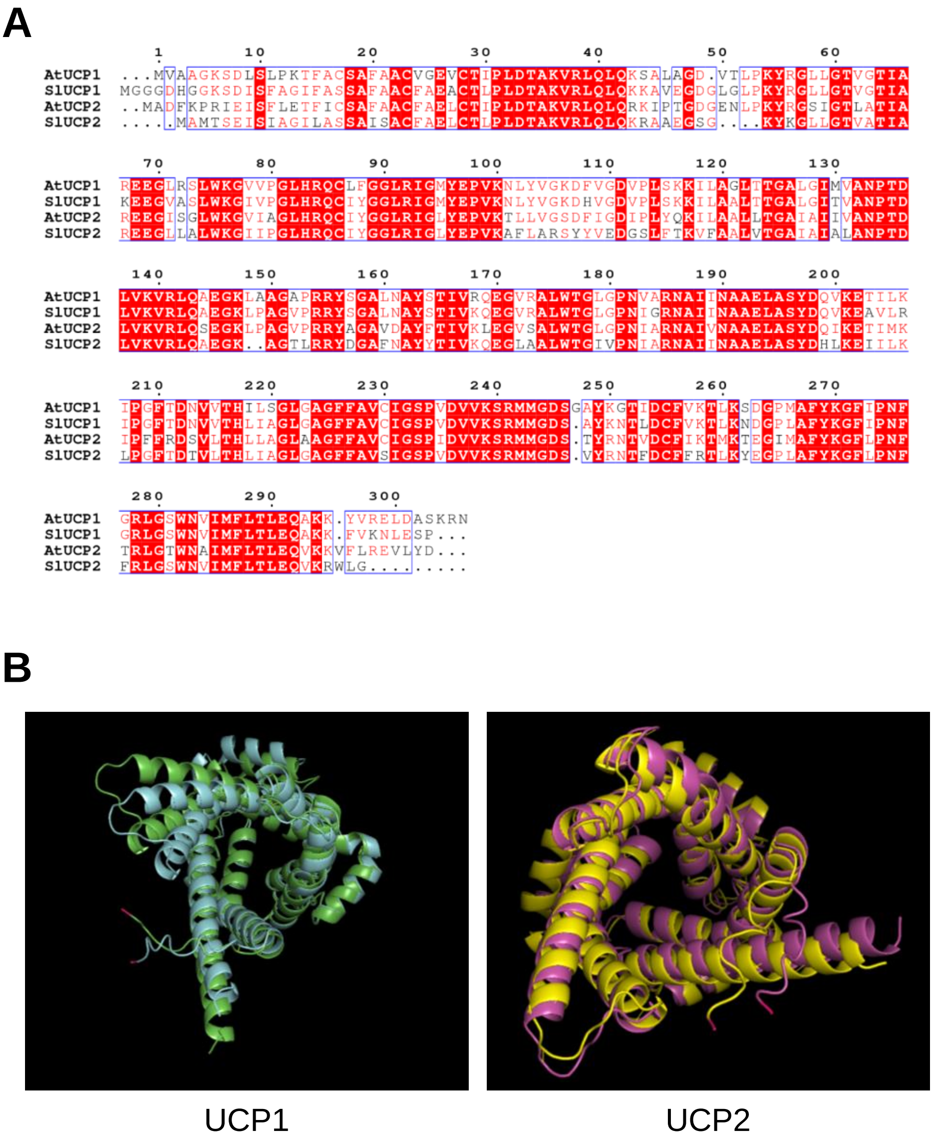


**Supplementary Figure 5.** Sequence and structural alignment of UCP1 and UCP2 from Arabidopsis and Tomato. **(A)** Multiple sequence alignment of Arabidopsis (At) UCP1 (AT3G54110, NP_190979.1; green) and UCP2 (AT5G58970, NP_568894.1; magenta) together with Tomato (Sl) UCP1 (Solyc09g011920, NP_001234584.1; blue) and UCP2 (Solyc09g031680, XP_004246961.1; yellow). AtUCP1 has 83.77% and 69.55% identity with SlUCP1 and SlUCP2 respectively. AtUCP2 has 70% and 71.05% identity with SlUCP1 and SlUCP2 respectively. (b) Alignments of the predicted structures for At and Sl UCP1 (left) and UCP2 (right).

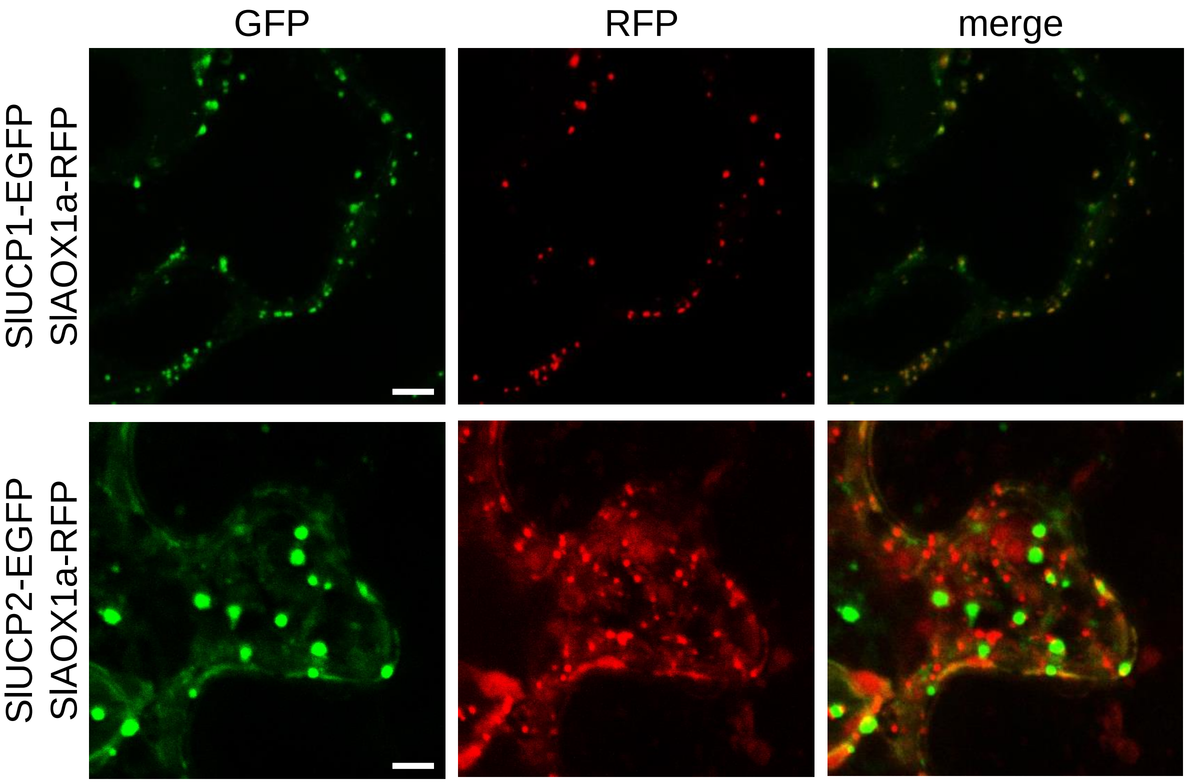


**Supplementary Figure 6.** Representative CLSM images of *Nicotiana benthamiana* leaves infiltrated with SlUCP1, SlUCP2 and SlAOX1a fused to fluorescent proteins. Confocal laser scanning microscope images were recorded from leaves 3 days after co-infiltration. Leaves were co-infiltrated with SlUCP1 or SlUCP2 fused to Enhanced Green Fluorescent Protein (EGFP) together with SlAOX1a fused to Red Fluorescent Protein (RFP). Both constructs are under the control of cauliflower mosaic virus 35S promoter. EGFP fluorescence: green, RFP fluorescence: red. Scale bars = 5 µm.
